# BoostMe accurately predicts DNA methylation values in whole-genome bisulfite sequencing of multiple human tissues

**DOI:** 10.1101/207506

**Authors:** Luli S. Zou, Michael R. Erdos, D. Leland Taylor, Peter S. Chines, Arushi Varshney, The McDonnell Genome Institute, Stephen C. J. Parker, Francis S. Collins, John P. Didion

## Abstract

Bisulfite sequencing is widely employed to study the role of DNA methylation in disease; however, the data suffer from biases due to coverage depth variability. Here we describe BoostMe, a method for imputing low quality DNA methylation estimates within whole-genome bisulfite sequencing (WGBS) data. BoostMe uses a gradient boosting algorithm, XGBoost, and leverages information from multiple samples for prediction. We find that BoostMe outperforms existing algorithms in speed and accuracy when applied to WGBS of human tissues. We also show that imputation improves concordance between WGBS and the MethylationEPIC array at low WGBS depth, suggesting improved WGBS accuracy after imputation.

## Background

DNA methylation is an epigenetic mark that is known to play a role in many fundamental biological processes, including differentiation, development, and gene regulation [1-2]. In mammals, DNA methylation occurs primarily on cytosines of CG dinucleotides (CpGs). CpG methylation marks convey epigenetic information across the lifespan, as they can be stably propagated through mitosis, and in special circumstances even through meiosis [3-7]. DNA methylation is an important mechanism for gene-environment interaction, and can thus influence health of cells, organs, and organisms.

DNA methylation is most commonly measured in cell lines or bulk tissue samples using microarrays or sequencing of bisulfite-converted DNA. These assays provide an estimate of the fraction of chromosomes in the cell population that are methylated at each CpG of interest (“beta” values). Microarrays such as the Illumina Infinium Methylation450k, and more recently the MethylationEPIC [8], are a cost-effective platform for measuring methylation in genes, promoters, and enhancers. However, the EPIC array measures only ~3% of all CpGs in the genome and has relatively little coverage of intergenic regions. In contrast, whole-genome bisulfite sequencing (WGBS) provides coverage of most of the ~28 million CpGs in the genome of an average tissue, giving it a clear advantage over the EPIC array. However, due to the high cost and sample input requirements of WGBS, it is often infeasible to generate deep-coverage data for a large number of replicates. Furthermore, methylation estimates from WGBS for CpGs sequenced at low depth are subject to error and are typically removed before performing downstream analysis [9]. One potential remedy for inefficiencies with WGBS is the generation of a small number of high-coverage reference samples in relevant tissues and disease states. These reference samples could be used to facilitate lower coverage and/or lower density methylation profiling in a larger number of samples. Such techniques have already been used to increase the power of GWAS studies by leveraging data from sparse yet cost-effective SNP arrays [10-12].

Machine and deep learning algorithms have shown promise in providing accurate beta value estimates after training on sparse data sets [13,14]. Prediction accuracy, however, is still far from the currently expected SNP imputation accuracy in GWAS [13], leading to the need for algorithm improvement. The most recent beta value imputation methods were based on either random forests [15] or deep neural networks [16]. A relatively new algorithm called extreme gradient boosting (XGBoost) has been shown to outperform both methods in accuracy and computational efficiency in data science competitions when highly predictive features can be constructed [17]. Previous imputation methods have also only classified beta values as fully unmethylated or methylated. This binarization of the data not only represents a loss of information but also ignores the possible significance of intermediate beta values as a conserved and biologically relevant genomic signature [18]. Specifically, although single chromsome methylation is binary, intermediate methylation in a population of cells has been shown to be a predominantly tissue specific signature that is enriched in genes, enhancers, and evolutionarily conserved regions [18,19]. Furthermore, although previous algorithms have constructed features that capture the local correlation structure of beta values [13,20] as well as information from the surrounding DNA sequence context [13,14,21-23], no algorithms have created features that incorporate information from multiple samples in the same tissue and/or disease state. This adaptation could improve prediction for CpGs that are not highly correlated to neighboring CpGs or strongly associated with their surrounding DNA context.

Importantly, machine and deep learning algorithms not only can impute missing values in sparse methylation data sets, but can also identify genomic features and sequence motifs associated with methylation patterns in different tissues [13,14,21,24-26]. For example, a previous random forest [13] algorithm applied to whole blood identified co-localized active transcription factor binding sites (TFBS), including those for ELF1, MAZ, MXI1, and RUNX3, to be predictive of beta values in whole blood. A recent deep learning algorithm [14] found that transcription factor motifs such as Foxa2 and Srf, which are both implicated in cell differentiation and embryonic development, were important to beta value prediction in mouse embryonic stem cells. These algorithms are therefore useful for characterizing methylation regulatory networks.

Methylation regulatory networks may have particular significance in complex diseases such as type 2 diabetes (T2D). The complexity of T2D is characterized by interactions between genetic and environmental factors acting in multiple tissues over time. Implicated tissues include pancreatic islets, skeletal muscle, adipose, liver, intestine, and brain. Genome-wide association studies (GWAS) have shown that the majority of T2D-associated loci lie in non-coding regions of the genome [27-29]. These loci therefore lack a clear relationship with any potential causal genes, underscoring the importance of identifying the epigenetic mechanisms by which they could affect gene expression.

In this work, we generated EPIC and WGBS data on 58 human samples from adipose, skeletal muscle, and pancreatic islets (**Additional file 1: Table S1**). Samples from adipose and skeletal muscle included those from patients with normal glucose tolerance (NGT) and T2D. We found 1) a high rate of missingness in the WGBS data and 2) discordance between WGBS and EPIC, biased towards low coverage and intermediate methylation sites. To address these issues, we developed an imputation method based on XGBoost called BoostMe, which is designed to leverage information from multiple independent samples from the same tissue type and disease state to impute low-coverage CpGs in WGBS data. We find that, for all tissues and all genomic contexts, BoostMe outperforms other methods, achieving the lowest error as well as the highest computational efficiency. We also examine the effect of imputation on WGBS accuracy by comparing raw WGBS and imputed beta values to those of the EPIC array. We find that discordance between EPIC and WGBS measurements at low WGBS depth is mitigated after imputation using BoostMe, supporting the use of imputation as an important preprocessing step for WGBS data analyses.

## Results and discussion

### Characterizing beta values in WGBS of adipose, skeletal muscle, and pancreatic islets

We generated WGBS and EPIC data from 58 samples of human adipose, skeletal muscle, and pancreatic islets. We discovered that, despite the relatively deep mean sequence coverage across samples (~30x genome-wide), there was a relatively high rate of missingness (CpG sequencing depth < 10x) (**Figure 1A**). The number of missing CpGs across all samples ranged from 2.6 million to 10.5 million, or roughly 10% to 40% of all ~25.5 million autosomal CpGs. We next explored the overlap between CpGs with a high rate of missingness and the underlying tissue-specific epigenomic architecture where they are located. We utilized previously published chromatin state segmentations for the corresponding tissues [30]. We found that missingness was spread across chromatin states (**Figure 1B**), with the highest raw numbers of missing beta values located in the Quiescent/Low Signal and Weak Transcription states (**Additional file 1: Figure S1**).

**Figure 1.**
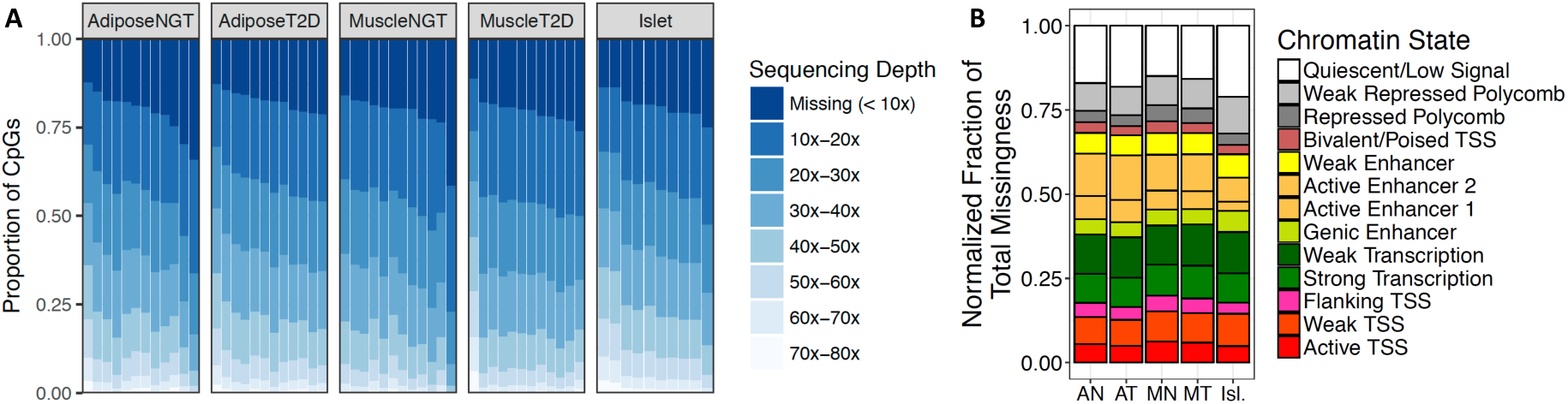
Characterization of WGBS missingness. **(A)** WGBS coverage (sequencing depth) distributions across tissue types and samples, visualized as the proportion of the ~25.5 million autosomal CpGs lying in each coverage interval. Each column is one sample from that tissue type. **(B)** WGBS missingness is distributed across chromatin states in all tissues. The normalized fraction of total missingness (y-axis) was calculated as the number of CpGs in each chromatin state that had missing beta values (sequencing depth < 10x) normalized by the total number of CpGs in that chromatin state for each tissue. Abbreviations: AN, adipose NGT; AT, adipose T2D; MN, muscle NGT; MT, muscle T2D; Isl., islets; NGT, normal glucose tolerance; T2D, type 2 diabetes.

Previous imputation work using array-based technology has shown that the beta value of a CpG is correlated with the beta values of its neighboring CpGs [13]. To determine the extent to which this may be true in WGBS, we quantified neighboring CpG similarity as a function of distance by calculating pairwise differences in methylation within chromatin states for each tissue and disease state combination (**Figure 2, Additional file 1: Figures S2, S3**). The majority (~70%) of CpG pairs genome-wide were highly similar, with an absolute difference in beta values less than 0.1 (**Figure 2A**). As distance between CpGs increased, chromatin states such as active and bivalent/poised transcription start site (TSS), strong and weak transcription, and quiescent/low signal had generally low differences, suggesting that neighboring beta values may be highly informative for prediction in these regions. In contrast, enhancer, flanking TSS, and repressed polycomb states exhibited larger differences as distance increased, suggesting that neighboring information alone may not be enough to make accurate predictions in these states, particularly when the nearest neighboring CpG is located at some distance.

**Figure 2.**
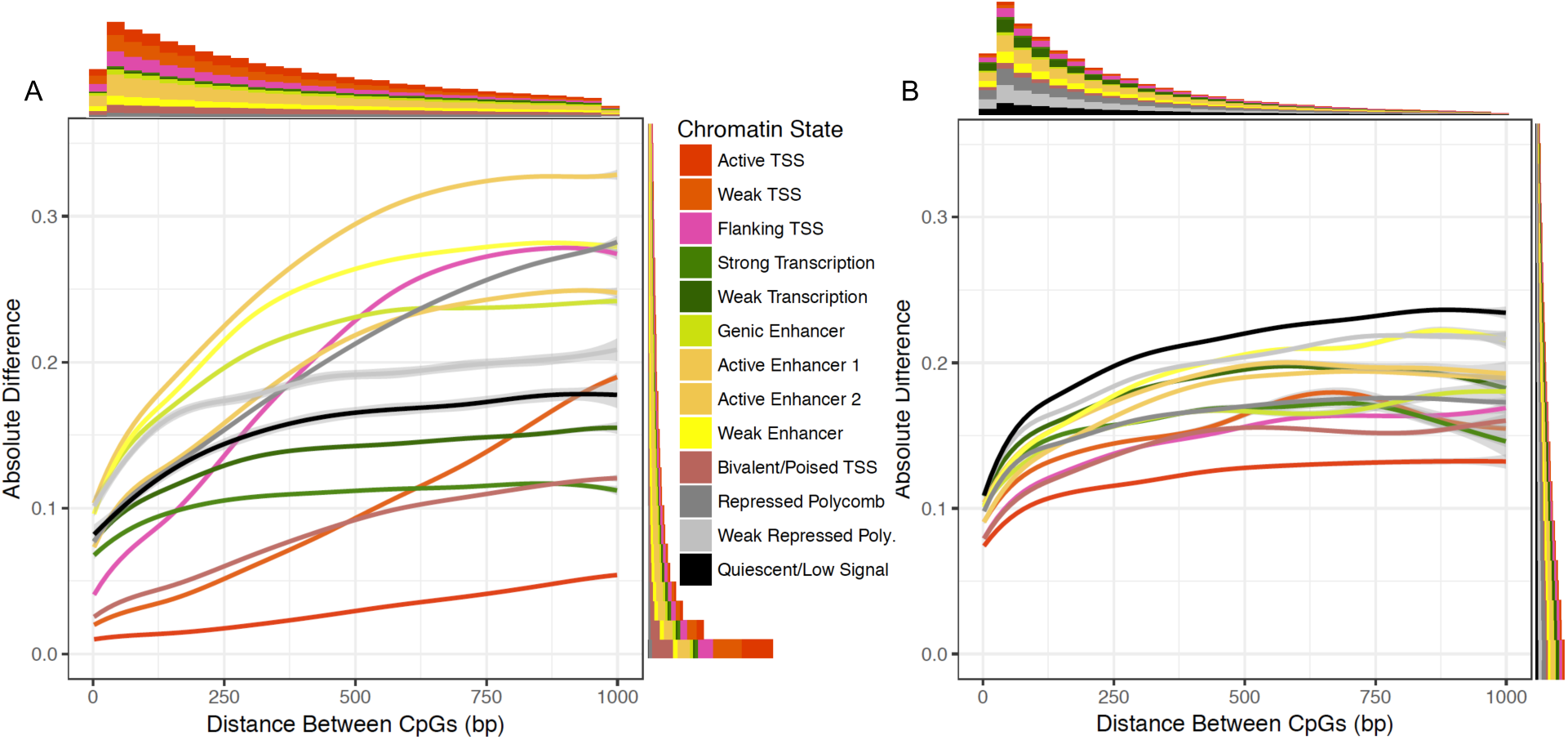
CpG methylation pairwise differences increase with distance and have different average behaviors across chromatin states. Differences were calculated using the average methylation value of each CpG across all 12 muscle NGT samples. Smoothed lines were fit using a generalized additive model. Average behaviors across chromatin states were similar across all tissue and disease state combinations examined in this work; see Additional file 1: Figures S2, S3. **(A)** Absolute pairwise differences within chromatin states genome-wide. We randomly sampled blocks of chromatin states genome-wide and used ~2 million pairwise differences for each chromatin state. Marginal histograms indicate the number of pairwise differences across chromatin states within the range of the graph. **(B)** Pairwise differences in blocks of consecutive CpGs with higher across-tissue variance. Variances were calculated across all 58 tissue samples; CpGs with methylation variances above the third quartile of variances were considered as having higher variances. Pairs of CpGs lying in the same high-variance region but different chromatin states were excluded from this analysis.

Since ~70-80% of CpGs are invariantly methylated across tissues and samples [31], we also calculated pairwise differences within regions of higher across-tissue variance. In contrast to the bimodal distribution of beta values genome-wide, average beta values in these high-variance blocks were more highly enriched for intermediate values (**Additional file 1: Figure S4**). Pairwise differences within these blocks exhibited less drastic changes over distance compared to the genome-wide analysis, and were more similar across chromatin states (**Figure 2B**). However, they had slightly larger magnitudes at low distances, where the bulk of differences occurred, indicating that even proximal CpGs may be less informative in these regions.

### BoostMe outperforms random forests and DeepCpG for methylation imputation

To address the high rate of missingness in our data, we developed BoostMe, a method for imputing beta values using WGBS data from at least three samples. Previous attempts at beta value imputation based on penalized functional regression [23], random forests [13], and deep neural networks [14] yielded relatively poor predictive accuracy genome-wide (RMSE > 0.23, AUROC < 0.93) (**Additional file 1: Table S2**). To improve on those methods, we implemented predictive models optimized for WGBS data using both random forest and gradient boosting [17] algorithms.

We constructed a total of 648 features designed to both parallel and improve upon previous work [13]. Prediction features constructed from the WGBS data included the nearest non-missing neighboring CpG beta values upstream and downstream of the CpG of interest, base-pair distance to the neighboring CpGs, and the average beta value of the CpG of interest in other samples from the same tissue and disease state (sample average). We also used tissue-specific reference data to create features that describe the genomic context of individual CpGs such as histone marks (n = 7), computational predictions of transcription factor binding sites (TFBSs) (n = 608), chromatin states (n = 13), and ATAC-Seq peaks (as a measure of DNA accessibility; see **Methods**, **Additional file 1: Table S3** for a full list of features).

Importantly, by testing the inclusion of different features in the model, we found that not all features had a beneficial effect on model accuracy (**Table 1**). The highest model accuracy was obtained when using the sample average, neighboring beta values and distances, ATAC-seq peaks, histone marks, GENCODE annotations, and chromatin states. Using these features, and after applying additional quality control exclusion criteria (**Methods**), the average number of CpGs usable for training and testing per sample was 20 million (range: 14.7 million - 21.2 million), and the average number of missing CpGs able to be imputed per sample (sequencing depth < 10x) was 2.6 million (range: 750,000 - 7.7 million).

**Table 1.** | Performance of BoostMe using different feature combinations.

| Features | RMSE<br>(all) | RMSE<br>(int.) | AUROC | AUPRC | Accuracy |
| --- | --- | --- | --- | --- | --- |
| Nearest non-missing upstream and downstream<br>neighboring beta values and distances (N) | 0.1515 | 0.2149 | 0.9489 | 0.9856 | 0.9370 |
| Sample average (A) | 0.09594 | 0.1430 | 0.9895 | 0.9942 | 0.9624 |
| A, N | 0.09330 | 0.1377 | 0.9902 | 0.9977 | 0.9639 |
| A, N, transcription factor binding sites | 0.09333 | 0.1378 | 0.9902 | 0.9977 | 0.9638 |
| A, N, recombination rate | 0.09330 | 0.1377 | 0.9902 | 0.9977 | 0.9639 |
| A, N, ATAC-seq peaks (P) | 0.09327 | 0.1377 | 0.9902 | 0.9977 | 0.9639 |
| A, N, histone marks (H) | 0.09322 | 0.1376 | 0.9902 | 0.9977 | 0.9639 |
| A, N, GENCODE annotations (G) | 0.09323 | 0.1376 | 0.9902 | 0.9977 | 0.9639 |
| A, N, chromatin states (C) | 0.09318 | 0.1375 | 0.9902 | 0.9977 | 0.9639 |
| A, N, P, H, G, C* | 0.09311 | 0.1373 | 0.9902 | 0.9977 | 0.9640 |

Performance was benchmarked on each sample by repeating the model training and testing process ten times using ten different random seeds. All metrics were calculated by averaging across all 58 samples. RMSE, root-mean-squared error; int., intermediate CpGs, defined as having a sample average methylation between 0.2 and 0.8; AUROC, area under the receiver operating characteristic curve; AUPRC, area under the precision-recall curve. Accuracy was calculated as the CpGs correctly predicted as methylated or unmethylated divided by the total number of CpGs. \*Final set of features used to benchmark performance.

We compared the performance of BoostMe and random forests with DeepCpG for predicting continuous beta values in WGBS data from adipose NGT (n = 12), adipose T2D (n = 12), muscle NGT (n = 12), muscle T2D (n = 12), and pancreatic islet (n = 10) tissue (**Figure 3A**). Due to memory limits, both BoostMe and random forests were trained on 1,000,000 randomly selected CpGs from a single sample, validated on a hold-out 500,000 CpGs, and tested on a hold-out set of 1,000,000 CpGs. We repeated this random sampling, training, and testing 10 times and averaged the results for each sample. DeepCpG was trained for each tissue and disease state combination as described in Angermueller *et al.* [14], using a total of ~10 million CpGs for training and ~5 million for validation. We evaluated DeepCpG models on a held-out random sample of 1,000,000 CpGs that also fit the BoostMe criteria for training and testing. We found that both BoostMe and random forests outperformed DeepCpG, achieving an average root-mean-squared error (RMSE) of 0.10, area under the receiver operating characteristic curve (AUROC) of 0.99, area under the precision-recall curve (AUPRC) of 0.99, and an accuracy of 0.96 (**Table 2**). Unlike previous methods [13,14], we trained on continuous beta values rather than binary values because of the available depth in our WGBS data, and found that this change improved overall RMSE by at least 0.06 and performed similarly for AUROC, AUPRC, and accuracy (**Additional file 1: Table S4**).

**Figure 3.**
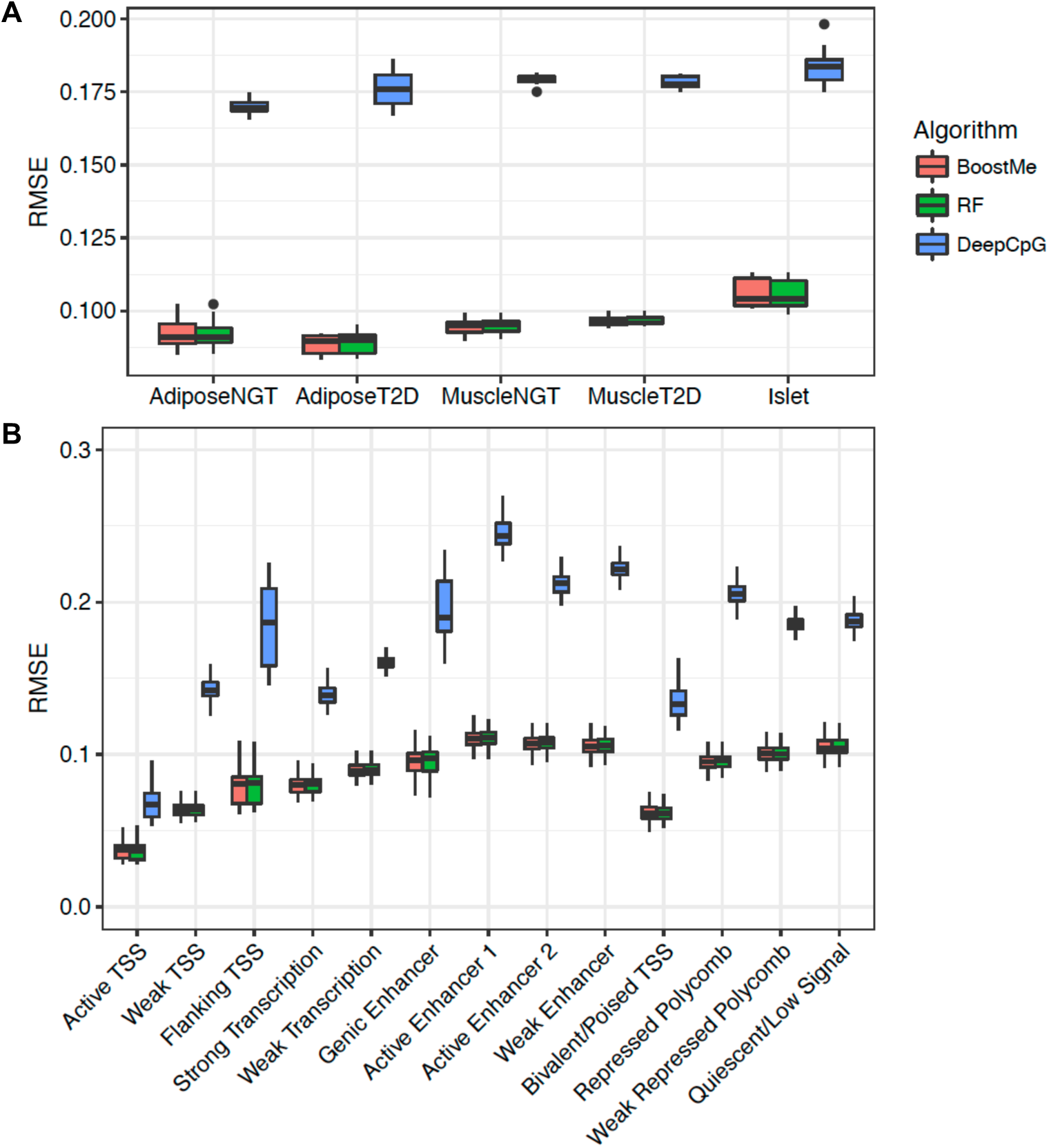
BoostMe and random forests outperform DeepCpG for predicting methylation values genome-wide. **(A)** Root-mean-squared error (RMSE) of BoostMe, random forests (RF) and DeepCpG for predicting methylation in all tissue and disease state combinations examined in this study. Data points represent performance on individual samples. NGT, normal glucose tolerance; T2D, type 2 diabetes. (B) RMSE of all algorithms by chromatin state.

**Table 2.** Genome-wide performance of different algorithms on predicting methylation values, averaged across tissues and samples.

| Algorithm | RMSE (all) | RMSE (int.) | AUROC | AUPRC | Accuracy | Resources | Time (hrs) |
| --- | --- | --- | --- | --- | --- | --- | --- |
| BoostMe | 0.09 | 0.13 | 0.99 | 0.99 | 0.96 | 16 CPUs | 0.33 |
| Random Forests | 0.09 | 0.13 | 0.99 | 0.99 | 0.96 | 16 CPUs | 14 |
| DeepCpG | 0.18 | 0.23 | 0.94 | 0.98 | 0.91 | 1 GPU | 144 |

RMSE, root-mean-squared error; int., intermediate CpGs, defined as having a sample average methylation between 0.2 and 0.8; AUROC, area under the receiver operating characteristic curve; AUPRC, area under the precision-recall curve.

To characterize performance patterns in different genomic contexts, we compared the performance of each algorithm within tissue-specific chromatin states (**Figure 3B**). Again, BoostMe and random forests outperformed DeepCpG in all chromatin states. In addition, all three algorithms exhibited the same trend across chromatin states, with the best predictive performance in TSS-associated states, which were strongly associated with low beta values (**Additional file 1: Figure S5**).

Beta values are bimodally distributed, with the majority of CpGs being either fully methylated or unmethylated; however, there is evidence that intermediate methylated CpGs are a conserved genomic signature that is often tissue-specific [18,19]. Furthermore, given our finding that regions of higher across tissue-variance tended to have average beta values in the intermediate range (**Additional file 1: Figure S4**), CpGs with intermediate average beta values may be more biologically significant, and therefore more important to predict accurately. Therefore, we also benchmarked all algorithms on intermediate beta values, defined as having a sample average methylation between 0.20 and 0.80 inclusive. We found similar trends in algorithm performance, with both BoostMe and random forests having an RMSE of 0.13 and DeepCpG an RMSE of 0.23 (**Table 2**).

We hypothesized that a uniform distribution of beta values in our training set would improve prediction at CpGs with intermediate beta values. We tested this by generating a training set using a biased sampling procedure that drew CpGs from each beta value decile with a frequency inversely proportion to its size. Contrary to our expectation, we found that this sampling procedure did not improve significantly the performance of BoostMe **(Additional file 1: Figure S6).**

We further examined BoostMe error as a function of distance to the nearest non-missing CpG across chromatin states, both genome-wide **(Figure 4A)** and within regions of higher across-tissue variance **(Figure 4B).** Trends in RMSE across distance strongly paralleled our previous analysis of pairwise differences in CpG methylation within chromatin states (**Figure 2**). As expected, the absolute prediction error was lowest for all chromatin states when there was a non-missing, neighboring CpG within 100 bp of the CpG of interest, which was true for the majority (~87%) of CpGs. The error increased for the smaller subset of CpGs where the nearest non-missing neighbor was farther away to varying degrees for each chromatin state: error in TSS states increased rapidly; transcribed states (strong transcription, weak transcription) remained relatively stable and low; and enhancer and inactive chromatin states had higher but generally stable error rates. Similar to the pairwise differences within regions of high across-tissue variance (**Figure 2B**), all chromatin states in these regions exhibited stably higher error rates, and had similar behaviors. Due to the small average block size for regions of high across-tissue variance (~533 bp on average), there was a lack of data past 200 bp which led to larger confidence intervals and less accurate smoothed line estimates.

**Figure 4.**
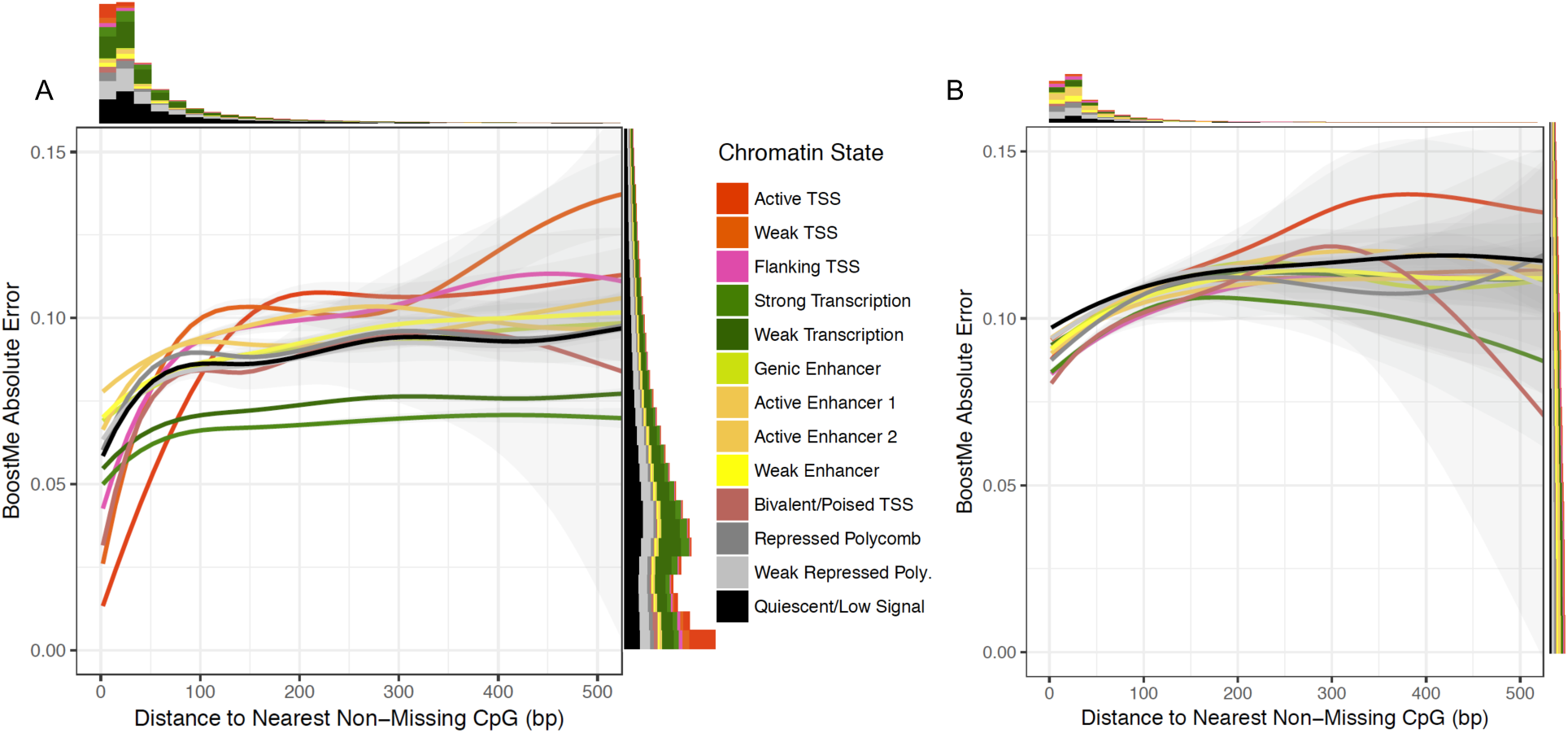
BoostMe error across chromatin states as a function of distance to the nearest non-missing CpG. Absolute prediction error was measured as the difference between the predicted and actual beta values for **(A)** a holdout test set of 5 million CpGs and **(B)** a holdout set of ~1.3 million CpGs in high-variance regions that met BoostMe criteria for training and testing. Gray shaded areas indicate confidence intervals for each smoothed line created using a generalized additive model. Marginal histograms display the number of pairwise differences across chromatin states within the range of the graph. Chromatin state corresponds to the chromatin state that the predicted CpG lied in.

Finally, we benchmarked the computational performance of all algorithms. BoostMe had a training runtime that was up to 40x faster than random forests using identical computational resources and up to 400x faster than DeepCpG (**Table 2**). Both BoostMe and random forests training times outperformed DeepCpG, which took multiple days to train due to the need to train CpG and DNA modules separately before training the joint module (**Methods**).

### Imputation reduces WGBS discordance with EPIC at low sequencing depth

To assess the effect of imputation on the quality of WGBS data, we first characterized the concordance of WGBS and EPIC array beta estimates at the same CpGs in the same samples (**Figure 5A**). As reported previously [32], WGBS and EPIC beta values were generally well-correlated (r^2^ = 0.92) (**Additional file 1: Figure S7**). However, we found that disagreement between the two platforms was concentrated at lower WGBS depth and intermediate beta values, with varying levels of discordance at high sequencing depth. Neither EPIC nor WGBS beta values can be considered the true methylation value of a particular CpG; however, since discordance between the two estimates was a function of sequencing depth, we hypothesized that discordance at low depth could be mostly attributed to WGBS inaccuracy, and discordance at high depth to EPIC inaccuracy.

**Figure 5.**
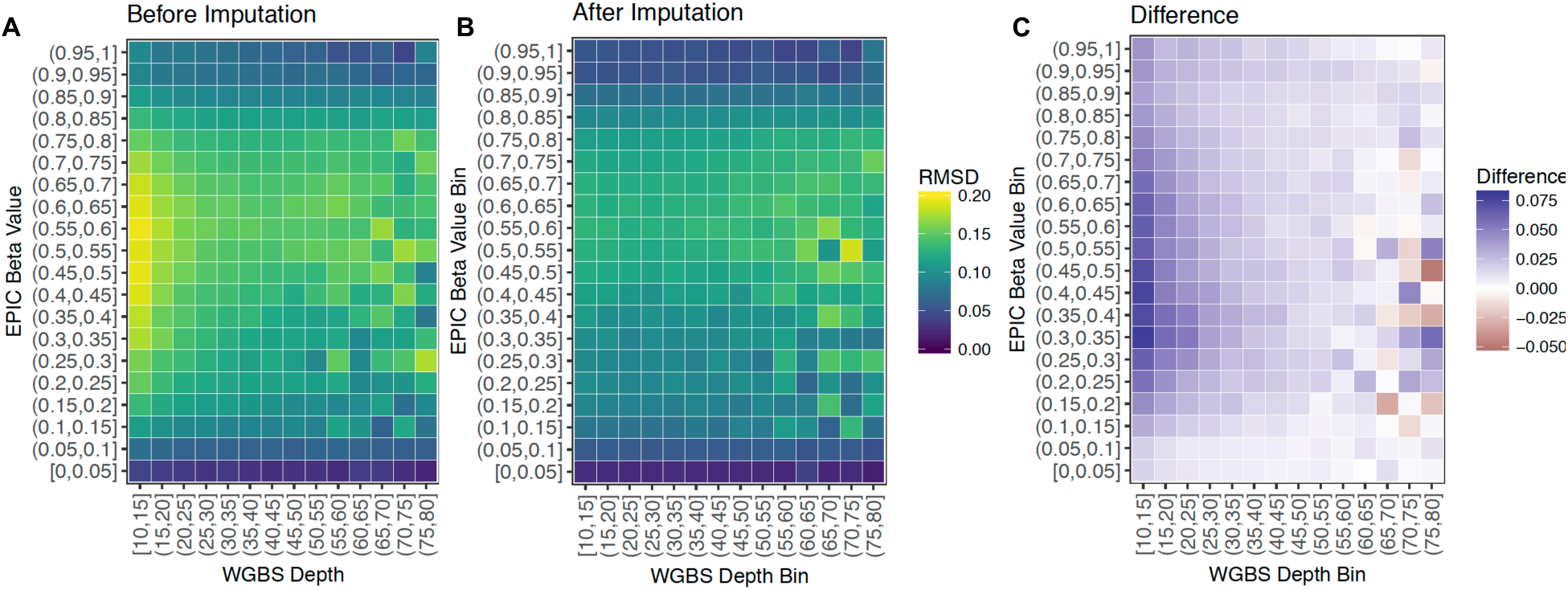
Imputation reduces discordance between WGBS and EPIC methylation estimates at low sequencing depth. **(A)** Root-mean squared discordance (RMSD) between EPIC and WGBS methylation estimates at CpGs common between the two platforms. X-axis: depth at which the CpG was sequenced, binned into intervals of 5. Y-axis: the beta value of the CpG as measured by the EPIC array, binned into intervals of 0.05. Yellow color indicates higher discordance. **(B)** RMSD between EPIC and imputed WGBS values at the same CpGs as in A. **(C)** Difference between A and B.

We then used BoostMe to impute and replace beta values common between the two platforms. We found that the discordance was mitigated (**Figure 5B**), particularly at lower WGBS depth and intermediate beta values (**Figure 5C**). Furthermore, we found that discordance mitigation at lower WGBS depth was robust with respect to the EPIC array probe type examined (**Additional file 1: Figure S8**). Discordance at higher depth was variable and in some cases increased after imputation (**Figure 5C**).

### BoostMe and random forests identify features important to general methylation levels

To interrogate differences in the methylation patterns of the different tissues and disease states, we examined the top variable importance scores output by random forests and BoostMe using all features (**Figure 6, Additional file 1: Figures S9, S10**).

**Figure 6.**
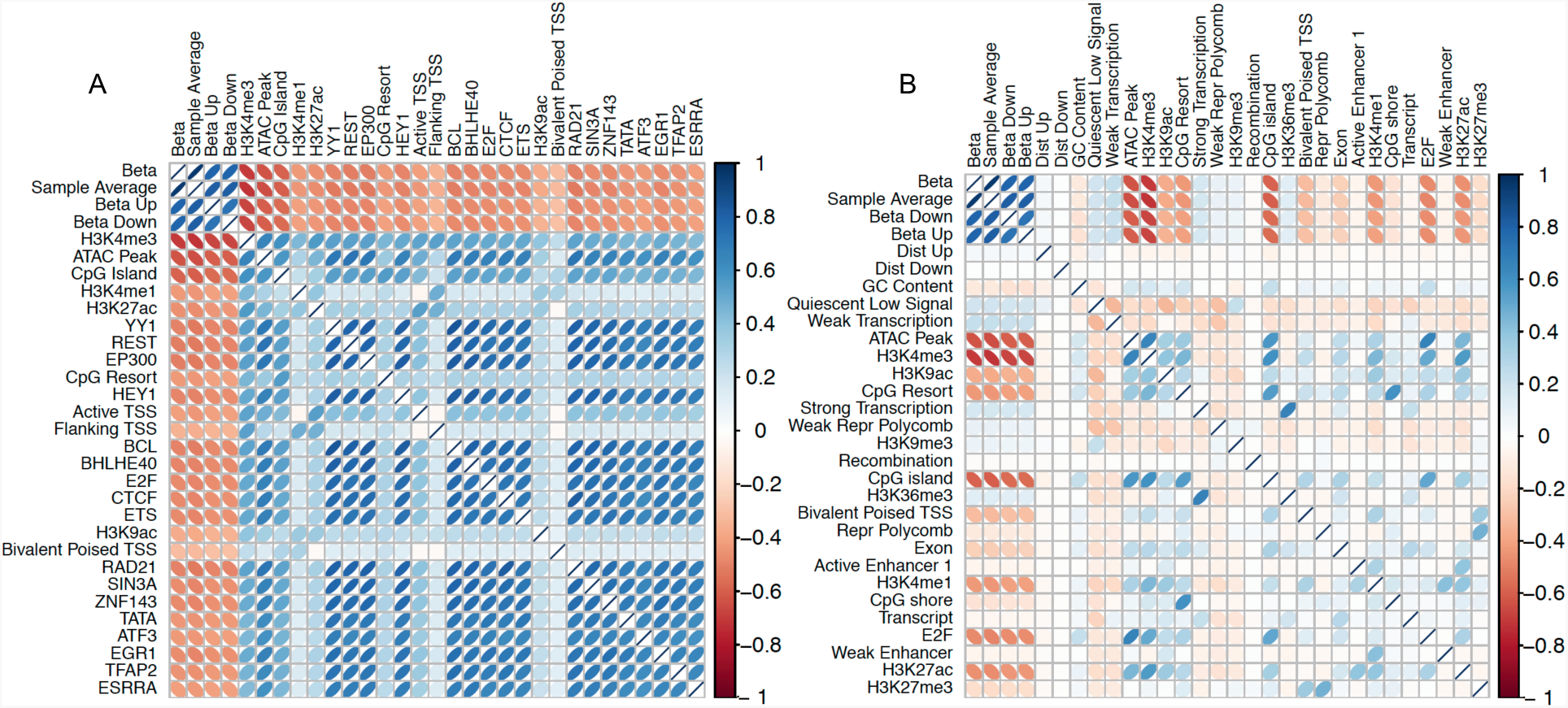
Random forests exhibit greater bias in favor of positively correlated features compared to BoostMe. Correlation among methylation beta value and top 30 features in descending order for adipose T2D as reported by **(A)** random forests and **(B)** BoostMe. Ranking was determined by aggregating the variable importance scores across 10 runs from all adipose T2D samples. Beta - beta value of the CpG of interest.

Both algorithms highly prioritized the sample average and neighboring CpG features, which were well-correlated with the beta value of the CpG of interest.

We found that random forests also ranked highly features that were negatively correlated with beta values, especially those associated with open chromatin and promoter regions such as H3K4me3, ATAC-Seq peaks, CpG islands, and the TSS chromatin states (**Figure 6A**). Random forests also identified several TFBSs previously shown to be methylation-sensitive such as YY1 [33,34], REST [35,36], and EP300 [37]. In concordance with previous results [13], we found that random forests were biased to rank highly features that are positively correlated with each other. This trend was particularly evident in the correlations among the top TFBSs identified, with all of them having some degree of overlap with each other (**Figure 6A**).

In contrast, BoostMe did not exhibit the same bias in favor of positively correlated features (**Figure 6B**). Since gradient boosting trees are trained sequentially, with each subsequent tree designed to reduce error from the previous tree, BoostMe is less likely to rank highly features that exhibit strong positive cross-correlations. Therefore, BoostMe more highly prioritized chromatin states that were positively correlated with methylation such as the quiescent/low signal and weak transcription chromatin states in addition to highly predictive features that were negatively correlated with methylation identified by random forests, such as ATAC-Seq peaks and H3K4me3. BoostMe did not report as many methylation-associated TFBSs as highly predictive, likely because of their high positive correlations with each other and with other features indicative of open chromatin, such as ATAC-Seq peaks and H3K4me3.

To further determine whether BoostMe or random forests could identify features that were important to specific tissues, we trained models using only TFBSs in tissue-specific regions of open chromatin as determined by ATAC-seq peaks. We found that the rankings were generally the same across tissues for both algorithms (**Additional file 1: Tables S5, S6**).

## Conclusions

Here we introduce BoostMe, a method for imputing low quality beta values within whole-genome bisulfite sequencing (WGBS) data. BoostMe is based on XGBoost, a computationally efficient gradient boosting algorithm that has seen widespread success in data science competitions [17]. Importantly, BoostMe leverages information from at least three samples and trains and predicts on continuous beta values. This framework allows BoostMe to outperform existing imputation methodology, including DeepCpG [13], a deep neural network method, in both speed and accuracy across tissues and genomic contexts. BoostMe also achieves lower RMSE than DeepCpG for intermediately methylated CpGs, which we found to be enriched in regions of high across-tissue methylation variance. Furthermore, using matched EPIC and WGBS data from the same samples, we have shown that BoostMe imputation reduces discordance between the two platforms, particularly at low WGBS depth. Overall, our results support the use of BoostMe as a preprocessing step to improve WGBS quality when multiple samples are available.

A notable limitation of BoostMe is its interpretability. Although it was previously reported that random forests identified TFBSs associated with methylation in whole blood [13], we found that neither BoostMe nor random forests identified noteworthy differences in variable importance scores between different tissues. On the other hand, DeepCpG, a deep neural network method, was able to identify differences in transcription factor motifs associated with prediction among the different tissues. For example, DeepCpG identified motifs of TFs important to a tissue type, such as EBF1 in adipose [38,39], ASCL2 in muscle [40], and FOXA1 in pancreatic islets [41,42], which have all been reported to be involved in regulating differentiation and development in their respective cell types. Thus, despite its relatively poor performance, DeepCpG may be superior for identifying tissue-specific differences. No algorithm examined in this work readily identified differences between NGT and T2D.

Similar to previous methodology [13,14], BoostMe relies on the locally correlated structure of neighboring CpGs to identify sample-specific differences. Although using neighboring information leads to an overall more accurate prediction for all algorithms examined in this work, the accuracy of this approach may not be robust for a subset of CpGs. To determine the local similarity of CpG methylation within WGBS and its effect on algorithm performance, we first calculated pairwise differences between beta values across chromatin states, finding that enhancer states generally had the largest differences, while TSS and transcribed states had the lowest. In concordance with this result, we found that all algorithms performed worst in active and weak enhancer chromatin states and best in TSS and transcribed states. Performance was slightly worse in regions of high across-tissue variance, where pairwise differences were generally larger.

Finally, to further characterize the relationship between informative neighboring CpGs and algorithm performance, we examined BoostMe error as a function of distance to the nearest non-missing CpG. The nearest non-missing beta values upstream and downstream were the second and third-most informative features in the model after the sample average. We found that error was lowest when the CpG of interest had a non-missing neighbor within 100 bp. This result parallels a previous study [43], which reported that methylation haplotype blocks, defined as areas of consecutive CpGs with r^2^ > 0.5, measure just 95 bp long on average. Although the majority (~87%) of CpGs did have a non-missing neighbor within 100 bp, the decreased performance for the remaining subset of CpGs may be a significant shortcoming of BoostMe and all neighbor-dependent imputation methods in general. Given these limitations, further work must be done to develop even more accurate imputation methodology that can identify sample-specific differences for prediction without depending heavily on informative neighboring CpGs, perhaps through incorporation of long-range interactions and other high-dimensional genomic and epigenomic features not considered in this work. Such methodology could potentially facilitate whole-genome imputation from a sparse subset of CpGs, with accuracy independent of neighboring CpG distance.

## Methods

### Sample collection

Muscle and adipose NGT and T2D samples were collected as previously described [4]. Briefly, we attempted to contact participants and participants’ relatives from previous diabetes-related studies [44-47] and also recruited subjects by newspaper advertisements. We excluded individuals with any diseases or drug treatments that might confound analyses. We defined glucose tolerance categories of NGT and T2D using World Health Organization (WHO) criteria [48]. Biopsies were performed by 9 experienced and well-trained physicians from 2009-2013 in 3 different study sites (Helsinki, Kuopio, and Savitaipale). The study was approved by the coordinating ethics committee of the Hospital District of Helsinki and Uusimaa. A written informed consent was obtained from all subjects.

Islet samples were collected as previously described [30]. Briefly, samples were procured from the Integrated Islet Distribution Program, the National Disease Research Interchange (NDRI), or ProdoLabs. Islets were shipped overnight from distribution centers, prewarmed in shipping media for 1-2 h before harvest, and cultured in tissue culture-treated flasks. Genomic DNA was then isolated from islet explant cultures and used for sequencing.

### Whole-genome sequencing

Whole genome sequencing libraries were generated from 50 ng genomic DNA fragmented by Covaris sonication. DNA end repair achieved using Lucigen DNA Terminator Repair Enzyme Mix. Sequencing adapters were added according to Illumina PE Sample Prep instruction. Libraries were size-selected on Invitrogen 4-12% polyacrylamide gels excising 200-250 bp fragments. Libraries were amplified with 10 PCR cycles and purified using AMPure beads (Beckman).

### Whole-genome bisulfite sequencing

Whole-genome bisulfite sequencing was performed using Epigenome/TruSeq DNA Methylation Kit (Illumina). Libraries were prepared for each sample using 50 ng of input DNA by denaturing the DNA at 98^°^C for 10 minutes. Bisulfite conversion was generated at 64^°^C for 2.5 h and DNA purified using EZ DNA Methylation Gold Kit (Zymo Research). Bisulfite converted libraries were generated by random-primed DNA synthesis, 3’ tagging, and purification using AMPure beads (Beckman). Sample-specific index sequences were added with 10 cycles of amplification.

Library quality was assessed using Qubit (Thermo Fisher Scientific) and Agilent Bioanalyzer. Paired-end 125bp sequencing was performed on Illumina HiSeq 2500 instruments to 30X genome coverage.

### EPIC array

Genomic DNA was extracted from each tissue using DNeasy Blood and Tissue Kits (QIAGEN), according to the manufacturer’s recommendations. 200ng of genomic DNA per sample was submitted to the Center for Inherited Disease Research at The Johns Hopkins University, where they were bisulfite-converted using EZ DNA methylation Kits (Zymo research), as part of the TruSeq DNA Methylation protocol (Illumina). DNA methylation was measured using the Illumina Infinium HD Methylation Assay with Infinium MethylationEPIC BeadChips according to manufacturer’s instructions.

### WGS data processing

Raw FASTQ files were evaluated with FastQC [49]. Adapter sequences were trimmed using Atropos [50], and reads with at least one pair shorter than 25 bp were excluded. Reads were aligned to the reference genome (GRCh37) using BWA MEM [51], followed by Samblaster [52] for marking duplicates.

### WGS variant calling

SNPs and indels were called separately for all sample BAM files using GATK HaplotypeCaller [53]. Variants were filtered using GATK Variant Quality Score Recalibration. Quality score cutoffs were chosen by comparing rates of discordance with SNP array genotypes.

### WGBS data processing

Raw FASTQ were pre-processed as above and aligned using bwa-meth [54]. Methylation values were extracted using the MethylDackel ‘extract’ command, including bias correct based on the values recommended by the ‘mbias’ command, and forward-and reverse-strand CpGs were merged with a minimum coverage cutoff of 10 (https://github.com/dpryan79/methyldackel). Methylation level data from the X and Y chromosomes were excluded.

### EPIC Array Data Processing

The EPIC data are part of a much larger, unpublished study. As such, all samples were processed jointly with other samples from the larger study. We processed raw signal idat files using minfi v1.20.2[55] with the Illumina normalization method. We analyzed the quality of each sample looking for outliers across a variety of measures including fraction of failed probes (detection p-value > 0.05), median methylated and un-methylated intensity, control probe signal (using the returnControlStat function from shinyMethyl v1.10.0 [56]), distribution of the overall methylation profile, and principle component analysis. None of the WGBS included in this study were flagged as outliers. In addition, we verified the identity of each sample by comparing genotypes assayed on the EPIC array to imputed genotypes using the HRC reference panel r1.1 [57] and Illumina Omni2.5 array genotypes.

For both the earlier 450k and recent EPIC Illumina methylation array, previous studies [58-61] have identified poor quality probes that either do not uniquely map to the reference genome or contain common genetic variation. These properties make the signal at these probes un-reliable. We removed such probes from the EPIC array. First, we removed cross-reactive probes on the EPIC chip by mapped non-control probes back to the entire bisulfite-converted genome, using Novoalign’s-b4 option, with allowance for up to three mismatches in the probe alignment (-R120 option). We kept only unique mapping probes. Second, we removed probes with a SNP within 10 bp of the 3’ end of the probe, within the target CpG itself, and finally, in the case of type I probes, if the variant overlaps the single base extension site. We used 10 bp as this cutoff is consistent with previous studies [59]. For SNPs we used common (MAF ≥ 1%) SNPs, indels or structural variation in the phase 3 1000 Genomes European dataset, common (MAF ≥ 1%) SNPs in the HRC reference panel r1.1, and SNPs appearing at all our own samples, even at low frequency, after imputation to the HRC reference panel. As a final step, we combined our blacklist with a previously published blacklist [61] for a total of 120,627 probes which were removed from analysis. In addition, we removed probes per tissue with a high detection p-value (p-value > 0.05 in ≥ 5% of samples from the larger study). After blacklist filters, we removed 578 adipose probes, 733 muscle probes, and 2,206 islet probes based on the per sample filters.

### Identification of higher across-tissue variance regions in WGBS

Using all 58 WGBS samples, we calculated the variance in methylation values for each CpG. We then searched for blocks of consecutive CpGs that had 1) variance above the third quartile of variance levels and 2) a non-missing methylation value in at least 20 samples, determined by looking at the distribution of variance values as a function of missing values (**Additional file 1: Figure S11**). This analysis identified approximately 200,000 blocks of high across-tissue variance CpGs genome-wide. Blocks contained an average of eight CpGs, spanned an average of 533 bp, and had higher relative enrichment in enhancer chromatin states (**Additional file 1: Figure S4**).

### Feature construction for BoostMe and random forests

We used the same 648 features in the BoostMe and random forest algorithms (see **Additional file 1: Table S2** for a detailed list). Prior to feature construction, we applied a further set of exclusion criteria to filter the CpGs included in training, validation, and testing. Only autosomal CpGs were used (n=25,586,776). We overlapped WGS data with the WGBS data from all samples and excluded CpGs for which the CG dinucleotide on either strand was disturbed by a SNP or indel that was 2 bp long. We also excluded all CpGs located in ENCODE blacklist regions [62].

#### CpG features

Features constructed from the WGBS data included neighboring CpG methylation values and the sample average feature. Neighboring CpG methylation values were taken within the sample of interest. For each neighbor, the methylation value as well as the base-pair distance from the neighbor to the CpG of interest were included as features. The sample average feature was created by taking the average of all samples within each tissue at the CpG of interest, not including the sample being interrogated. Samples in which the CpG was not sequenced above 10x coverage were excluded from the calculation. CpGs without a measurement above 10x coverage from at least two additional samples were also excluded.

#### Genomic features

We constructed both general and tissue-specific genomic features. General genomic features were the same across all tissues and included GC content, recombination rate, GENCODE annotations, and CpG island (CGI) information. GC content data was downloaded from the raw data used to encode the gc5Base track on hg19 from the UCSC Genome Browser [63,64]. DNA recombination rate annotations from HapMap were downloaded from the UCSC hg19 annotation database(http://hgdownload.soe.ucsc.edu/goldenPath/hg19/database/). CpG island coordinates were obtained from UCSC browser. CpG island shores and shelves were calculated from CpG island coordinates by taking 2 kb flanking regions. GENCODE v25 transcript annotations were downloaded from the GENCODE data portal (ftp://ftp.sanger.ac.uk/pub/gencode/Gencode_human/release_25).

Tissue-specific genomic features included ATAC-seq, chromatin states, histone marks, and transcription factor binding sites (TFBS). These features were all binary, with 0 indicating that the CpG of interest did not overlap that feature, and 1 indicating overlap. Chromatin state annotations were obtained from a previously published 13 chromatin state model for 31 diverse tissues that included islets, skeletal muscle, and adipose [30]. This model was generated from cell/tissue ChIP-seq data for H3K27ac, H3K27me3, H3K36me3, H3K4me1, and H3K4me3, and input from a diverse set of publicly available data [19, 65-67]. ATAC-seq data was obtained from previously published studies for islets [30], skeletal muscle [68], and adipose [69]. TFBS data was obtained as described in [68], with additional PWMs from [70]. TFBS data was filtered for each tissue by the ATAC-seq feature to only include hits overlapping an ATAC-seq peak. We merged hits from multiple motifs of the same transcription factor to reduce the number of variables included in the algorithm and optimize computational efficiency.

### BoostMe and random forests implementation

For BoostMe we used the xgboost package (version 0.6-4) [17] in R [71] (version 3.3.1). For random forests we used the ranger package (version 0.6.0) in R, which facilitates random forest training and testing on multiple CPUs [72]. In the final algorithm, we set num_trees to 500 to balance computational time and accuracy, and used default values for other parameters after finding that performance was robust to different settings.

For both algorithms, we used regression trees to predict a continuous methylation value between 0 and 1 for CpGs of interest. Algorithms were trained on individual samples within each tissue and disease state combination. We trained only on CpGs with at least 10x coverage and no more than 80x coverage. Random forest variable importance was calculated using the mean decrease in variance at each split as implemented in the ranger package. BoostMe variable importance was evaluated for each variable as the loss reduction after each split using that variable as implemented in the xgboost package.

Due to memory limits, algorithms were trained on a random sample of 1,000,000 CpGs from all available CpGs within a sample, validated on a hold-out set of 500,000 CpGs, and benchmarked on another hold-out set of 1,000,000 CpGs. To account for possible biases that would arise from only training on a small subset of the over 20 million CpGs available for training from each sample, we repeated the process of randomly sampling CpGs for training, validation, and testing ten times for each sample using ten different random seeds and averaged the results.

### DeepCpG implementation

We implemented DeepCpG (version 1.0.4) as described in Angermueller *et al.* (2017) [14]. Briefly, for each of the five tissue and T2D status combinations (adipose NGT, adipose T2D, muscle NGT, muscle T2D, and islet) the data was first divided by chromosome into training (chr. 1, 3, 5, 7, 9, 11, 13, 15), validation (chr. 16, 17, 18, 19, 20, 21, 22), and test sets, corresponding to a rough 40-20-40 split. The DNA module and CpG module were trained on separate NVIDIA Tesla K80 GPUs and the performance of each module was evaluated individually on the test set. The joint module was trained with the best-performing DNA and CpG modules, and its predictions were used for final benchmarking. In contrast to original single-cell bisulfite implementation of DeepCpG which was trained and tested on binary methylation values, we trained and tested on continuous methylation values to parallel our implementation of BoostMe and random forests. We find that this change made no difference in the accuracy of the model (**Additional file 1: Table S4**).

We experimented with six different hyperparameter combinations for each DNA model, including three architectures (CnnL2h128, CnnL2h256, CnnL3h256) and two dropout rates (0, 0.2). We then selected the best-performing combination based on AUC and reported the motifs significantly matching the filters from the first convolutional layer of that model [73]. Similarly, we tested both RnnL1 and RnnL2 for the CpG model for each tissue. For the joint module, we tested JointL1h512, JointL2h512, and JointL3h512. The best-performing joint model was selected to evaluate RMSE, AUC, AUPRC, and accuracy for each tissue. We used a default learning rate of 0.001 for all models. Similar to random forests and BoostMe, performance was generally robust with respect to different architectures. For a detailed explanation of all model architectures, see http://deepcpg.readthedocs.io/.

## Declarations

### Ethics approval and consent to participate

The study was approved by the coordinating ethics committee of the Hospital District of Helsinki and Uusimaa. A written informed consent was obtained from all subjects.

### Consent for publication

Not applicable

### Availability of data and materials

The datasets analyzed from the current study will be submitted upon publication. We will also release BoostMe as an R package through Github and Bioconductor upon publication.

### Author’s contributions

JPD conceived of the study. LSZ developed the software and ran experiments. LSZ, JPD, and FSC wrote the manuscript. MRE and MGI generated the data. DLT, PSC, AV, and SCJP contributed to data analysis. All authors read and approved the final manuscript.

## Acknowledgements

The authors would like to thank members of the Collins laboratory for helpful discussions and Christof Angermueller for his assistance with DeepCpG. This manuscript is dedicated to Peter Chines.

